# Efficient search for informational cores in complex systems: Application to brain networks

**DOI:** 10.1101/2020.04.06.027441

**Authors:** Jun Kitazono, Ryota Kanai, Masafumi Oizumi

**Affiliations:** Department of General Systems Studies, Graduate School of Arts and Sciences, The University of Tokyo, Meguro-ku, Tokyo, Japan; Araya, Inc., Minato-ku, Tokyo, Japan

## Abstract

To understand the nature of the complex behavior of the brain, one important step is to identify “cores” in the brain network, where neurons or brain areas strongly interact with each other. Cores can be considered as essential sub-networks for brain functions. In the last few decades, an information-theoretic approach to identifying cores has been developed. In this approach, many-to-many nonlinear interactions between parts are measured by an information loss function, which quantifies how much information would be lost if interactions between parts were removed. Then, a core called a “complex” is defined as a subsystem wherein the amount of information loss is locally maximal. Although identifying complexes can be a novel and useful approach to revealing essential properties of the brain network, its practical application is hindered by the fact that computation time grows exponentially with system size. Here we propose a fast and exact algorithm for finding complexes, called Hierarchical Partitioning for Complex search (HPC). HPC finds complexes by hierarchically partitioning systems to narrow down candidates for complexes. The computation time of HPC is polynomial, which is dramatically smaller than exponential. We prove that HPC is exact when an information loss function satisfies a mathematical property, monotonicity. We show that mutual information is one such information loss function. We also show that a broad class of submodular functions can be considered as such information loss functions, indicating the expandability of our framework to the class. In simulations, we show that HPC can find complexes in large systems (up to several hundred) in a practical amount of time when mutual information is used as an information loss function. Finally, we demonstrate the use of HPC in electrocorticogram recordings from monkeys. HPC revealed temporally stable and characteristic complexes, indicating that it can be reliably utilized to characterize brain networks.

**Author summary:** An important step in understanding the nature of the brain is to identify “cores” in the brain network, which can be considered as essential areas for brain functions and cognition. In the last few decades, a novel definition of cores has been developed, which takes account of many-to-many interactions among elements of the network. Although considering many-to-many interactions can be important in understanding the complex brain network, identifying cores in large systems has been impossible because of the extremely large computational costs required. Here, we propose a fast and exact algorithm for finding cores. We show that the proposed algorithm enables us to find cores in large systems consisting of several hundred elements in a practical amount of time. We applied our algorithm to electrocorticogram recordings from a monkey that monitored electrical activity of the brain with electrodes placed directly on the brain surface, and demonstrated that there are stable and characteristic core structures in the brain network. This result indicates that our algorithm can be reliably applied to uncovering the essential network structures of the brain.

## 1 Introduction

The brain achieves its highly sophisticated cognitive functions through the interaction of many neurons. To understand the neural mechanisms of brain functions, it is important to uncover network structures in the brain [1–5]. An effective way to characterize these network structures is to identify “cores” of the network where neurons strongly interact with each other. The cores can be considered as the most important sub-networks for various functions and cognition (see [4, Chapter 6] for a detailed review). In the literature, cores have been defined in many ways using different approaches, such as maximal cliques [6], k-cores [7–10], and rich-clubs [9–11]. Most of these methods are based on graph representations of systems, wherein the dependence between two nodes is represented as the weight of the edge connecting the two nodes. Although this graph-based analysis is easy to use, it does have a limitation: since a graph is fully described by one-to-one relations between nodes, graph-based analysis inevitably omits the effects of many-to-many interactions; and although these are not predicted by one-to-one interactions alone, they are still important for understanding complex interactions in the brain.

To overcome the limitations of graph-based analysis, we utilize an information-theoretic approach [12, 13] to cores that takes account of many-to-many interactions. In this approach, the degree of interactions between parts in a system is measured by an information-theoretic measure, such as mutual information. More specifically, it is quantified by how much information would be lost if a system were cut into parts and interactions between the parts were removed. Then, roughly speaking, a core is defined as a subsystem wherein the amount of information loss is larger than that in any of its supersets. We call such cores defined this way “informational cores” or “complexes” [12, 13].

The original idea of the complex was proposed in the Integrated Information Theory (IIT) of consciousness [12–17]. IIT hypothesizes that complexes in the brain correspond to the loci of consciousness, i.e., the areas where consciousness arises. Nevertheless, we can utilize complexes not only to study consciousness but also to analyze other systems unrelated to consciousness. This is because the complex is based on the information theory and can therefore be applied to any stochastic system, in principle at least.

Despite the general applicability of the complex to stochastic systems and its potential superiority to other graph-based approaches, only a few studies have utilized complexes to analyze systems, and the sizes of the systems analyzed were small [18–20]. This is because searching for complexes is extremely difficult in large systems: the search computation time grows exponentially with the number of elements in the system, and identifying complexes is virtually impossible even in systems with only dozens of elements.

In this paper, we propose a fast and exact algorithm, which we call “Hierarchical Partitioning for Complex search” (HPC). HPC searches for complexes by hierarchically dividing subsystems. HPC is exact when the measure of the information loss satisfies a certain mathematical property, i.e., monotonicity. Mutual information satisfies this property and is therefore used in this study. We also show that a class of submodular functions, where “submodular” is a mathematical property of set functions [21, 22], satisfies this monotonicity, indicating the expandability of our framework to the class of functions. HPC can identify complexes in polynomial time. More specifically, HPC enables us to find complexes of a system consisting of several hundreds of elements, which in turn makes possible the search for complexes in, for example, multi-channel EEG and ECoG, which typically involve several hundreds of electrodes, in a practical amount of time.

The rest of the paper is organized as follows. In Section 2.1, we outline important concepts that are needed to define the complex. In Section 2.2, we explain information loss, in particular the mutual information. In Section 2.3, we explain how the information loss is utilized to quantify the strength of interactions in a system. In Section 2.4, we define the complex. Then, in Section 2.5, we introduce our new HPC algorithm. In Section 2.6, we show the relation between monotonicity and submodularity. In experiments in Section 3, we first demonstrate how HPC works by taking a simple model as an example. Second, we evaluate the computation time of HPC. Third, we demonstrate how the algorithm can be applied to real neural data using an open ECoG dataset as an example [23]. Finally, in Section 4, we discuss some limitations of HPC and prospects of this study.

The MATLAB codes of HPC are available at https://github.com/oizumi-lab/PhiToolbox.

## 2 Methods

### 2.1 Outline of important concepts

“Complex” is a rather complicated concept to understand. Before we give a formal mathematical definition of complexes, we first outline two important concepts, information loss functions and minimum information partitions, which are needed to define complexes, and then outline complexes.

#### Information loss function

Information loss functions measure the strength of dependence between parts from an information-theoretic viewpoint. Information loss functions quantify how much information is lost if a subsystem is “cut” into two smaller parts, where “cut” means removing the interactions between the two parts. If the two parts are independent of each other, no information is lost while if the parts are strongly dependent on each other, much information is lost. As we detail in the next subsection, mutual information and integrated information in IIT can be interpreted as information loss functions.

#### Minimum information partition (MIP)

The amount of information loss depends on how a subsystem is partitioned. As an example, consider a subsystem {1, 2, 3, 4} consisting of the two independent parts {1, 2} and {3, 4} (Fig 1). If the subsystem is vertically cut into the two parts {1, 2} and {3, 4} as shown in Fig 1A, no information is lost. In contrast, if the subsystem is partitioned differently, as shown in Fig 1B, some amount of information is lost. The minimum information partition (MIP) of the subsystem is that partition among all possible partitions for which the information loss is minimum [12, 13, 15]. In this example, the vertical cut in Fig 1A is the MIP. Thus, the MIP cuts the system with its weakest link.

**Fig 1.**
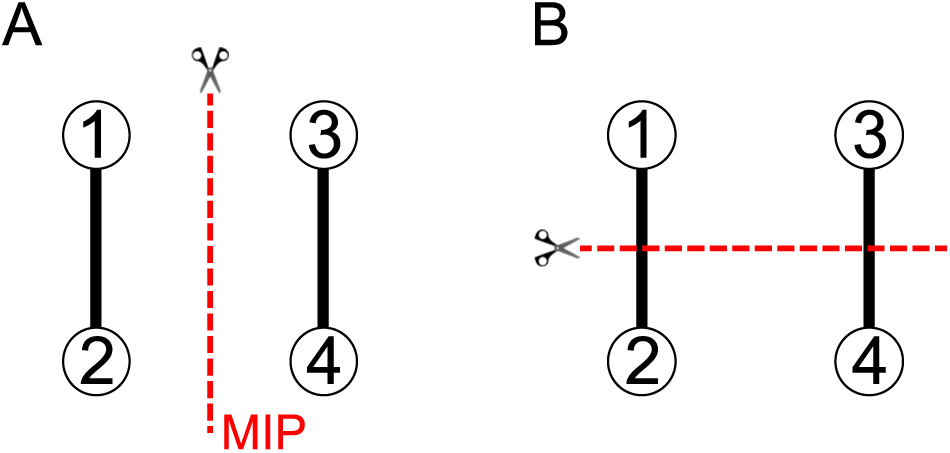
Schematic of Minimum Information Partition (MIP). We consider a subsystem consisting of four elements {1, 2, 3, 4}. The nodes connected by edges are dependent on each other, while others are not. If the subsystem is vertically cut into the two parts {1, 2} and {3, 4} as shown in Fig 1A, no information is lost. In contrast, if the subsystem is partitioned horizontally as shown in Fig 1B, some amount of information is lost. In this case, the vertical cut in Fig 1A is the MIP.

#### Complex

Complexes are defined by using the concept of MIP: a complex is a subsystem such that the amount of information loss when it is partitioned with its MIP is larger than those of all its supersets [12, 13]. If we add elements to a complex, the amount of information loss of the extended subsystem for its MIP is smaller than that of the complex. Intuitively speaking, a complex is more strongly unified than any of its supersets, and adding elements to a complex inevitably includes a weaker link.

### 2.2 Information loss function

In this subsection, we explain the definition of information loss functions. Information loss functions measure the amount of information loss when interactions between parts are removed. After introducing a general definition of information loss functions, we consider the mutual information as an example of information loss functions. As we describe in Section 2.5, the mutual information satisfies essential properties for the proposed algorithm for complex search. For this reason, we use the mutual information as an information loss function in this paper.

We consider a probabilistic system consisting of *N* elements with a distribution *p*(***x***_*V*_) = *p*(*x*_1_, …, *x*_*N*_). Here, *V* denotes the set of indices (*V* = {1, …, *N*}) and ***x***_*V*_ denotes (*x*_1_, …, *x*_*N*_) = (*x*_*i*_)_*i*∈*V*_. Similarly, for a subset *S* ⊆ *V*, ***x***_*S*_ denotes (*x*_*i*_)_*i*∈*S*_. For example in the case of a multi-agent system, each variable *x*_*i*_ can be a state of an agent, and *N* can be the number of agents. In the case of an EEG measurement of brain activity, *x*_*i*_ can be a signal from an electrode and *N* can be the number of electrodes.

We consider a bi-partition (*S*_L_, *S*_R_) of a subset *S* that partitions *S* into two disjoint parts, *S*_L_ and *S*_R_ (*S*_R_ = *S*\*S*_L_). We then consider a “disconnected” distribution, 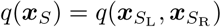, where some types of interaction between *S*_L_ and *S*_R_ are removed. We define the amount of information loss caused by the partitioning, *f* (*S*_L_; *S*_R_), as the Kullback-Leibler divergence between the original distribution *p* and the disconnected distribution *q*:

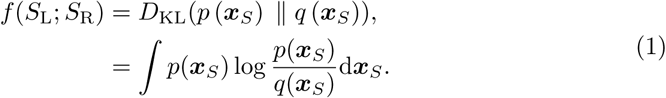

The Kullback-Leibler divergence, in general, can be interpreted as information loss when *q* is used to approximate *p* [24]. Thus, the information loss function *f* can be interpreted as information loss caused by removing interactions between parts.

Since *S*_L_ is the complement of *S*_R_ in *S*, determining *S*_L_ automatically specifies *S*_R_. Therefore, an information loss function *f* (*S*_L_; *S*_R_) can be considered as a set function of *S*_L_ whose domain is the powerset of *S*. To emphasize this, we introduce the following notation:

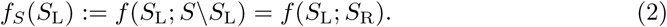

There are various types of information loss function, depending on what kind of “interactions” are removed [25–29]. An example is integrated information, which was originally introduced in IIT [13, 15, 16]. Several variants have also proposed [25–28, 30, 31]. Integrated information can be interpreted as information loss when “causal” interactions between subsystems are removed [25]. Integrated information becomes large when two parts strongly affect each other across different time points (see [25–27, 30, 31] for more details).

Another example of an information loss function is mutual information, which we use in this study. Mutual information quantifies statistical dependence between parts. Let us consider the following disconnected probability distribution *q*,

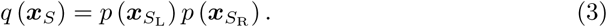

In this model *q*, the two parts *S*_L_ and *S*_R_ are independent, which means any kind of dependence between *S*_L_ and *S*_R_ is completely removed. Then, by substituting Eq (3) into Eq (1), the mutual information *I*(*S*_L_; *S*_R_) between the two parts *S*_L_ and *S*_R_ is given by

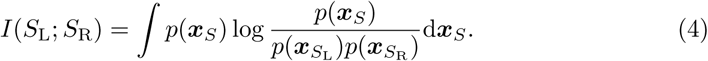

The mutual information measures how strongly two subsystems are statistically dependent on each other. It becomes large when the two parts are strongly dependent, and becomes 0 when the two parts are independent; that is, 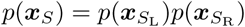. The mutual information Eq (4) is also represented as

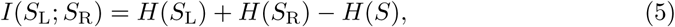

where *H*(·) represents the entropy, e.g., *H*(*S*) = − ∫*p*(***x***_*S*_) log *p*(***x***_*S*_)d***x***_*S*_. Equivalently, we can rewrite the above equation as a set function of *S*_L_:

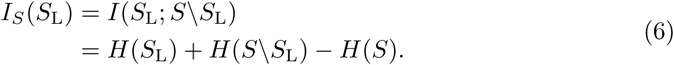

We use Eqs. (5) and (6) in Sections 2.3 and 2.6.

The mutual information has two important mathematical properties which are utilized in this study, as shown later. The first is symmetric-submodularity, which enables a fast and exact search for minimum information partitions (MIPs) [32, 33] (Section 2.3). The second is monotonicity, which is the basis of our new algorithm HPC (Section 2.5).

### 2.3 Minimum information partition

In this subsection, we introduce the Minimum Information Partition (MIP) and an efficient algorithm for searching for MIPs, which is called Queyranne’s algorithm.

#### 2.3.1 Definition of MIPs

The MIP is the bi-partition for which the amount of information loss is minimum among all bi-partitions. Mathematically, the MIP 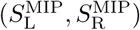 of a subsystem *S* is defined as follows:

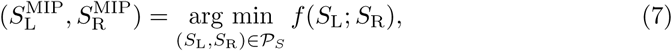

where 𝒫_*S*_ denotes the set of all the bi-partitions of the subset *S*. Since the information loss function *f* (*S*_L_; *S*_R_) is represented as a set function *f*_*S*_ (*S*_L_), the search for the MIP can be equivalently formulated as a minimization problem of the set function *f*_*S*_ (*S*_L_):

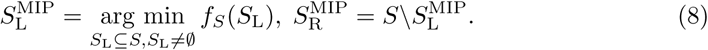

We represent the amount of information loss for the MIP as

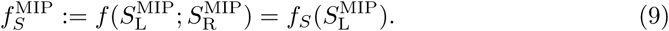

Similarly, we represent the mutual information for the MIP as 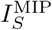. Hereinafter, we refer to a bi-partition as a partition, for simplicity.

By 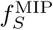, we can evaluate the “irreducibility” of the subsystem *S*. When 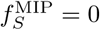, this means the subsystem *S* consists of independent parts, i.e., the subsystem *S* can be reduced to independent parts. In contrast, when 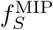 is nonzero, the subsystem *S* cannot be reduced to independent parts. No matter how the subsystem *S* is cut into parts, at least as much information as 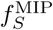 is lost.

#### 2.3.2 Algorithm for searching for MIPs

If we search for MIPs by exhaustively comparing all the partitions, the computation time grows exponentially with the number of elements in the system. In previous studies, we utilized a mathematical concept, submodularity, to reduce the computation time [32–34]. In particular, we used a submodular-based algorithm called Queyranne’s algorithm. In the next paragraphs, we first introduce the definition of submodularity and then introduce Queyranne’s algorithm.

##### Submodularity

Submodularity is a property of set functions that is analogous to the concavity of continuous functions. Specifically, submodularity is defined as follows:

###### Definition 1

(Submodularity). *A set function f* : 2^*S*^ → ℝ *is submodular if it satisfies the following inequality for any A, B* ⊆ *S:*

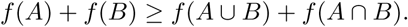

The mutual information *I*_*S*_ (*S*_L_) is submodular as a function of *S*_L_, and the entropy *H* is also submodular. The submodularity of the mutual information can be easily derived from the submodularity of the entropy by using Eq (6) [22].

If a submodular function *f* : 2^*V*^ → ℝ satisfies *f* (*S*) = *f* (*V*\*S*) for any subset *S* ⊆ *V*, the function *f* is called symmetric-submodular function. The mutual information *I*_*S*_ (*S*_L_) is a symmetric-submodular function defined over the powerset of *S*.

In general, given any submodular function *f* : 2^*S*^ → ℝ, a function *g*_*S*_ : 2^*S*^ → ℝ defined as

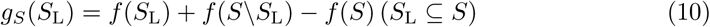

is symmetric-submodular [22]. In this paper, we call this type of symmetric-submodular functions “symmetrized” submodular functions. The mutual information, which is symmetric-submodular, is especially “symmetrized”-submodular. We use the concept of symmetrized submodular functions in Section 2.6.

##### Queyranne’s algorithm

If a set function *f* : 2^*S*^ → ℝ is symmetric-submodular, we can exactly and efficiently find the minimum of the function by Queyranne’s algorithm [35]. Thus, we can use Queyranne’s algorithm to find MIPs when the mutual information is used as an information loss function [32, 34]. The computation time of the algorithm is *O*(|*S*|^3^), where *S* indicates the number of elements of |*S*|. This is much smaller than an exhaustive search, wherein the computation time is *O*(2^|*S*|^).

### 2.4 Complex: informational core of a system

In this subsection, we introduce the definition of a complex [12, 13]. We also introduce a main complex, which is a stronger definition of a complex [12, 13].

A subsystem is called a complex if the amount of information loss for its MIP is nonzero and larger than those of all its supersets. The mathematical definition of a complex is given as follows.

#### Definition 2

(Complex). *A subset S* ⊆ *V is called a complex if it satisfies* 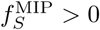 *and* 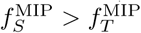 *for any of its superset T (T* ⊃ *S and T* ⊆ *V)*.

A schematic explanation of the definition of a complex is shown in Fig 2. The subsystem {3, 4, 5} is a complex if it has greater *f* ^MIP^ than all its supersets; that is, 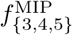 is larger than 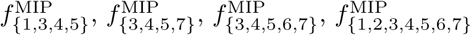, and so on.

**Fig 2.**
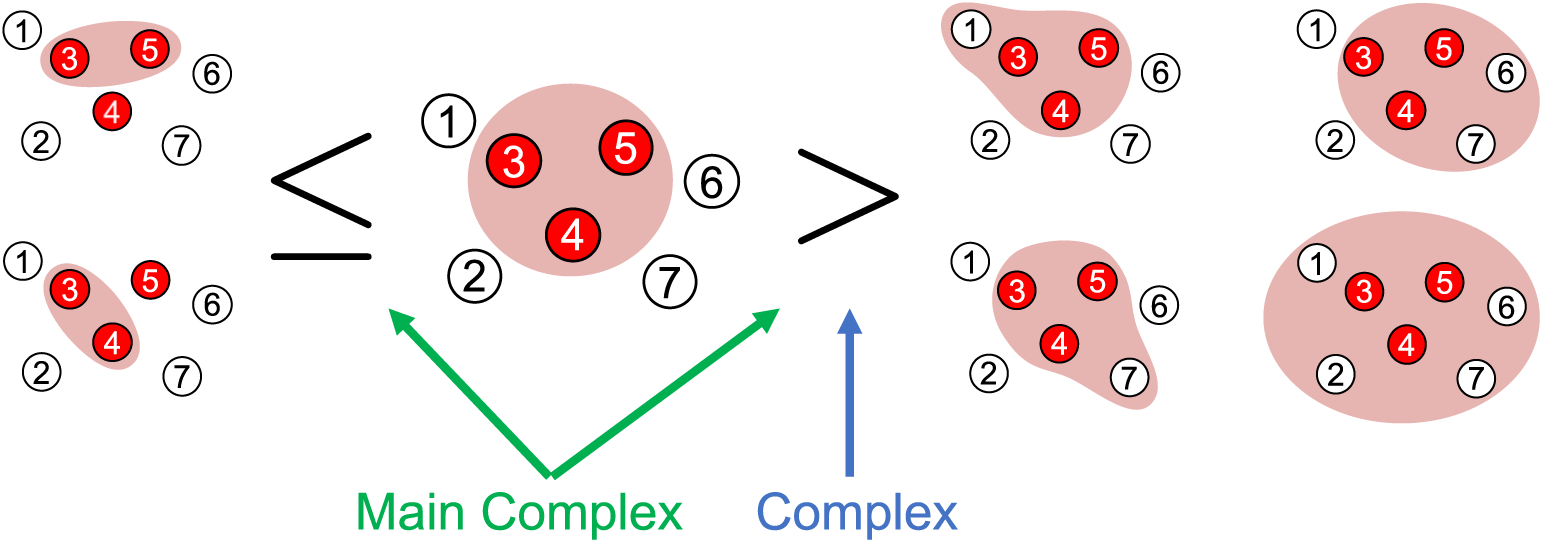
Schematic of the definitions of complex and main complex (Definitions 2 and 3). A subsystem *S* is a complex, if *S* has larger *f* ^MIP^ than any supersets of *S*. In this example, the subsystem {3, 4, 5} is a complex if 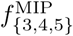 is larger than 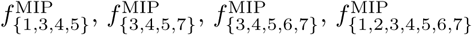, and so on. The subsystem *S* is a main complex if *S* is a complex, and *S* has a larger *f* ^MIP^ than any subsets of *S*. The subsystem {3, 4, 5} is a main complex if {3, 4, 5} is a complex and 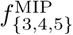 is larger than 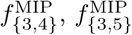, and 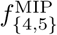.

The whole system *V* is a complex if it satisfies 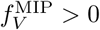 by definition. We define *f* ^MIP^ = 0 for single elements because we cannot consider partitions of a single element. Therefore, single elements cannot be complexes.

A subsystem is called a main complex if the amount of information loss for its MIP is larger than those of all its supersets, and is also larger than or equal to those of its subsets. In other words, a complex is called a main complex if the amount of information loss with its MIP is larger than or equal to those of all its subsets.

#### Definition 3

(Main complex). *A complex is called a main complex if it satisfies* 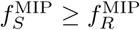 *for any of its subset R (R* ⊂ *S)*.

A schematic explanation of the definition of main complexes is shown in Fig 2. The subsystem {3, 4, 5} is a main complex if it has *f* ^MIP^ larger than those of all its supersets, and also has *f* ^MIP^ equal to or larger than those of all its subsets, i.e., {3, 4}, {3, 5}, and {4, 5}.

A complex can be regarded as an informational core in the sense that when elements are added to it, its irreducibility always decreases, as quantified by *f* ^MIP^. A main complex can be also regarded as an informational core but in a stronger sense, in that its irreducibility always decreases, both when elements are added to it and also when they are removed from it. Thus, a main complex can be considered as a locally-most-irreducible subsystem.

We end this subsection by mentioning a property of main complexes which will be utilized in HPC. A main complex does not include smaller complexes by definition of main complexes and complexes. Conversely, if a complex *S* does not include smaller complexes, then *S* is a main complex. This can be easily shown by contradiction. Thus, a main complex is a complex that does not include smaller complexes. This property of main complexes is stated as a lemma below.

#### Lemma 4.

*A complex is a main complex if and only if the complex does not include other complexes*.

### 2.5 Searching for complexes

If we search for (main) complexes by brute force, we need to find the MIPs of all the *O*(2^*N*^) subsets, and then compare the *f* ^MIP^ of the subsets to check if each subset satisfies the definition of a (main) complex (Defs. 2 and 3). In contrast, by using Hierarchical Partitioning for Complex search (HPC), we need to find the MIPs of only *N* − 1 subsets.

In what follows, we describe the algorithm HPC. We first explain that the mutual information satisfies an important mathematical property, i.e., monotonicity. We then show that, by taking advantage of the monotonicity of the mutual information, HPC enables us to find complexes efficiently.

#### 2.5.1 Monotonicity of mutual information

The mutual information has a well-known mathematical property, i.e., monotonicity, as described below. Let us consider the mutual information between *A* and *B, I*(*A*; *B*), where *A* and *B* are sets of elements. Then, if we add another set of elements *C* to *A*, the mutual information does not decrease. That is,

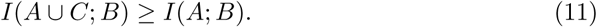

Also if we add *C* to *B, I*(*A*; *B* ∪ *C*) ≥ *I*(*A*; *B*). This inequality means that the mutual information monotonically increases as elements are added. We refer to this mathematical property as “monotonicity.” The above inequality is equivalent to the non-negativity of the conditional mutual information

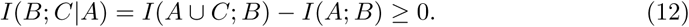

To utilize this concept in HPC, we restate the monotonicity in the context of partitioning a system. Let *T* be a subset of *V* and (*T*_L_, *T*_R_) be a partition of *T*. For any non-empty subset *S*_L_ of *T*_L_ (*S*_L_ ⊂ *T*_L_), the following inequality holds:

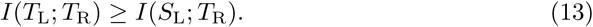

#### 2.5.2 Auxiliary theorems

We show several auxiliary theorems that are the basis of our new algorithm HPC.

By using the monotonicity of the mutual information Eq (13), we can show an inequality that particularly holds for the mutual information for MIPs, which is denoted by *I*^MIP^, as follows.

##### Proposition 5.

*Let T be a subset of V, and* (*T*_L_, *T*_R_) *be the MIP of T*. *If a subset S of T (S* ⊂ *T) intersects both T*_L_ *and T*_R_ *(S* ∩ *T*_L_ ≠ ø *and S* ∩ *T*_R_ *≠* ø*), then* 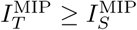.

*Proof*. We show 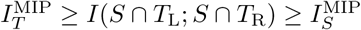. The first inequality immediately follows from the monotonicity of the mutual information Eq (13):

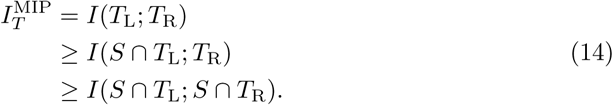

Then, the second inequality holds by definition of the MIP: Since (*S* ∩ *T*_L_, *S* ∩ *T*_R_) is a partition of *S*, that is, (*S* ∩ *T*_L_) ∪ (*S* ∩ *T*_R_) = *S* and (*S* ∩ *T*_L_) ∩ (*S* ∩ *T*_R_) = ø, the mutual information *I*(*S* ∩ *T*_L_; *S* ∩ *T*_R_) is always larger than or equal to 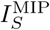. □

A schematic explanation of (the proof of) Proposition 5 is shown in Fig 3. In this example, a system *T* = {1, 2, 3, 4, 5, 6, 7} is partitioned into *T*_L_ = {1, 2, 3, 4} and *T*_R_ = {5, 6, 7} with its MIP. A subsystem *S* = {2, 4, 5, 7} is a subset of *T* (*S* ⊂ *T*). Then, 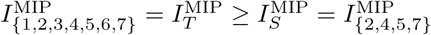, because *S* intersects both *T*_L_ and *T*_R_; that is, *S* ∩ *T*_L_ = {2, 4, 5, 7} ∩ {1, 2, 3, 4} = {2, 4} *≠* ø and *S* ∩ *T*_R_ = {2, 4, 5, 7} ∩ {5, 6, 7} = {5, 7} *≠* ø.

**Fig 3.**
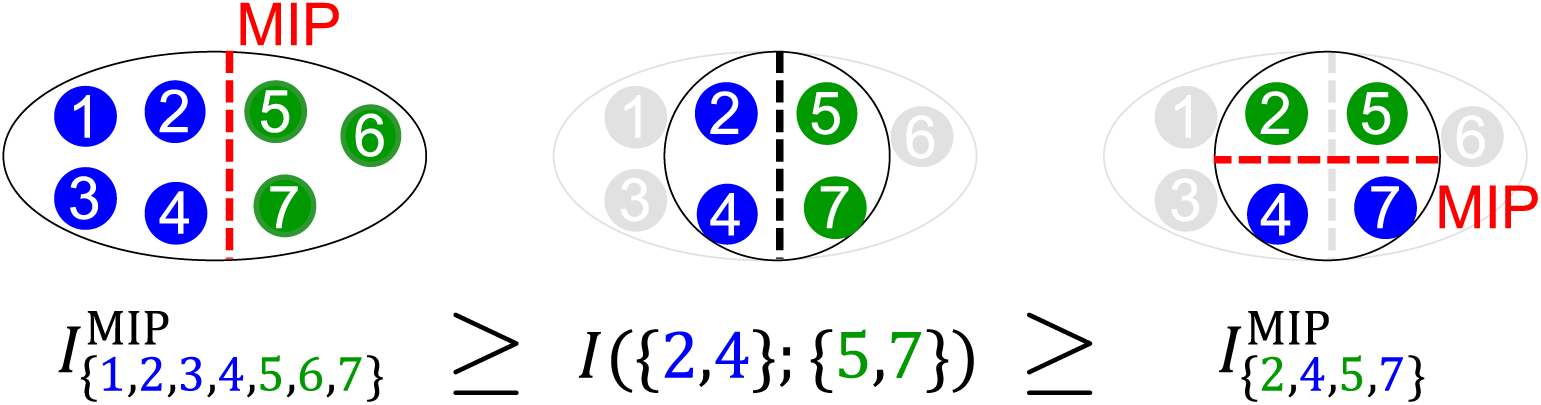
Schematic explanation of the proof of Proposition 5. Here we consider a system *T* = {1, 2, 3, 4, 5, 6, 7}. This system *T* is partitioned by its MIP into two subsystems *T*_L_ = {1, 2, 3, 4} and *T*_R_ = {5, 6, 7}. The boundary of the MIP of *T* is indicated by a red-dashed vertical line. Then, we consider a subsystem *S* = {2, 4, 5, 7} ⊂ *T*. Since *S* intersects both *T*_L_ and *T*_R_ (*S* ∩ *T*_L_ = {2, 4} *≠* ø and *S* ∩ *T*_R_ = {5, 7} *≠* ø), the first inequality 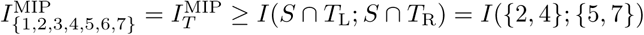 holds. The second inequality 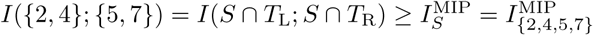 holds by definition of the MIP. The boundary of the MIP of *S* is indicated by a red-dashed horizontal line.

The inequality in Proposition 5 means that the subset *S* has smaller *I*^MIP^ than that of its superset *T*, and therefore *S* does not satisfy the definition of complexes (Def. 2). Thus, the lemma below follows immediately.

##### Lemma 6.

*Let T be a subset of V, and* (*T*_L_, *T*_R_) *be the MIP of T*. *If a subset S of T (S* ⊂ *T) intersects both T*_L_ *and T*_R_ *(S* ∩ *T*_L_ ≠ ø *and S* ∩ *T*_R_ ≠ ø*), then S is not a complex*.

Lemma 6 means that a proper subset *S* of *T* (*S* ⊂ *T*) must be included in *T*_L_ or *T*_R_ to be a complex. In other words, any complex that is a subset of *T* is included in *T*_L_ or *T*_R_. Therefore, the following proposition holds.

##### Proposition 7.

*Let T be a subset of V, and* (*T*_L_, *T*_R_) *be the MIP of T*. *Then, any complex S that is a subset of T (S* ⊆ *T) satisfies one of the following mutually exclusive conditions (a)–(c)*.

a. *S* = *T*,
b. *S* ⊆ *T*_L_,
c. *S* ⊆ *T*_R_.

Proposition 7 is the main basis of our algorithm, Hierarchical Partitioning for Complex search (HPC), as will be shown in Section 2.5.3.

#### 2.5.3 Hierarchical Partitioning for Complex search

In the following paragraphs, we explain our algorithm, Hierarchical Partitioning for Complex search (HPC). HPC primarily consists of two steps. The first step is listing candidates of (main) complexes. HPC narrows down candidates for (main) complexes by hierarchically partitioning a system. The second step is screening the candidates to find (main) complexes. In the succeeding paragraphs, we explain these two steps.

##### Hierarchical partitioning for listing candidates of complexes

Hierarchical Partitioning for Complex search (HPC) is schematically shown in Fig 4, and a pseudo-code is given in Algorithm 1. HPC enables the finding of all the complexes and main complexes by hierarchical partitioning of the system with the minimum information partitions.

**Fig 4.**
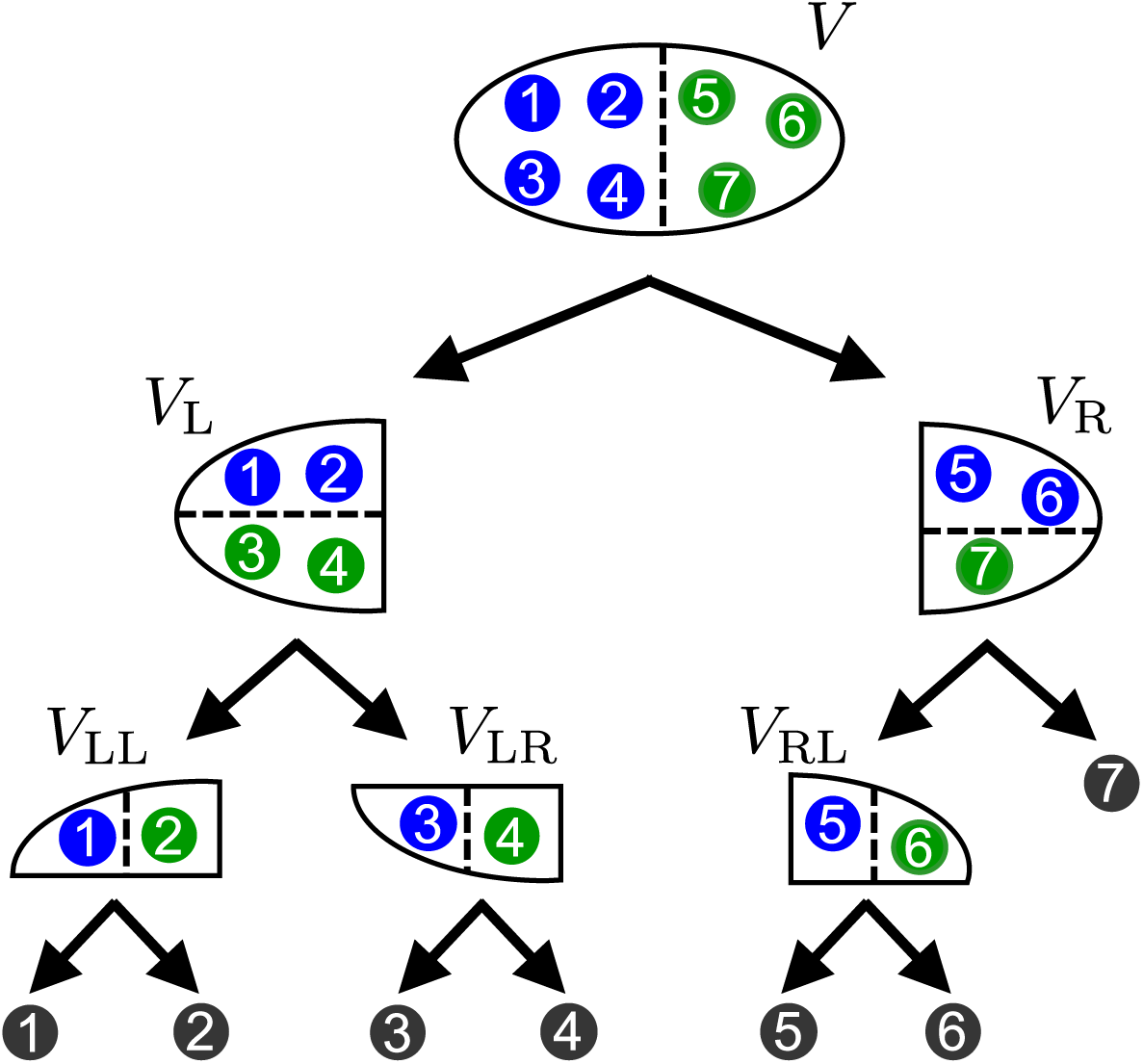
Schematic of Hierarchical Partitioning for Complex search (HPC). In HPC, a system is hierarchically partitioned by MIPs until the system is decomposed into single elements. In this example, the whole system *V* = {1, 2, 3, 4, 5, 6, 7} is divided by its MIP (indicated by a dashed line) into *V*_L_ = {1, 2, 3, 4} and *V*_R_ = {5, 6, 7}. Then, *V*_L_ is divided into *V*_LL_ and *V*_LR_, and *V*_R_ into *V*_RL_ and {7}. Finally, the whole system *V* is decomposed into seven single elements.

As is shown in Fig 4, HPC starts by dividing the whole system with its MIP, and then repeatedly divides the subsystems with their MIPs until the whole system is completely decomposed into single elements. This procedure in HPC is summarized as follows:

1. Find the MIP (*V*_L_, *V*_R_) of the whole system *V* and divide the whole system *V* into two subsystems *V*_L_ and *V*_R_.
2. Find the MIPs of the subsystems found in the previous step, *V*_L_ and *V*_R_, and divide them into (*V*_LL_, *V*_LR_) and (*V*_RL_, *V*_RR_), respectively.
3. Repeat this division until the whole system is decomposed into single elements.

After the procedure above, we obtain the set of hierarchically partitioned subsystems, i.e., *V, V*_L_, *V*_R_, *V*_LL_, *V*_LR_, *V*_RL_, *V*_RR_, and so on. We consider all the set of subsystems 𝒱 = {*V, V*_L_, *V*_R_, *V*_LL_, *V*_LR_, *V*_RL_, *V*_RR_, …}, excluding single elements. Then, the following theorem holds.

###### Theorem 8.

*Any complex S* ⊆ *V belongs to* 𝒱 *(S* ∈ *V)*.

*Proof*. By repeatedly applying Proposition 7 to *V, V*_L_, *V*_R_, *V*_LL_, *V*_LR_, *V*_RL_, *V*_RR_, and so on, we obtain the desired result. □

Thus from Theorem 8, 𝒱 can be seen as the set of candidates of complexes. Also, Theorem 8 means that complexes in a system form a “nested” hierarchy: if there are two complexes, one includes the other, or the two are disjoint. They never partially overlap each other. This nested hierarchy of complexes cannot be derived only from the definition of complexes (Def. 2), because the definition does not specify the relation between complexes. For the nested hierarchy to hold, the inequality of the mutual information for MIPs (Proposition 5) is necessary.

As described above, HPC decomposes the whole system into *N* single elements, by dividing a subsystem into two subsystems at every step. This means that HPC divides the system *N* − 1 times. That is, HPC evaluates MIPs of *N* − 1 subsets. This number is much smaller than the number of subsets evaluated in the exhaustive search, 2^*N*^ − *N* − 1, which is the number of subsets consisting of more than one element.

Next, we describe the way to select complexes and main complexes from the set of candidates 𝒱.

##### Selection of complexes from the candidates

After the hierarchical partitioning procedure described above, we need to check whether each candidate of complexes belonging to 𝒱 is actually a complex or not in accordance with Def. 2. We can efficiently check this by taking advantage of the hierarchical (tree) structure. Please see Part A in S1 Text for details.

##### Selection of main complexes from the candidates

As stated in Lemma 4, a main complex is a complex that does not include smaller complexes. Thus, a main complex is a locally farthest complex from the root (the whole system) in the tree. Based on this, we can easily find main complexes. See Part B in S1 Text for details.

#### 2.5.4 Time complexity of HPC

We derive a rough estimate of the total time complexity for searching for complexes by HPC. Here we only consider the first step of HPC (hierarchical partitioning for listing candidates of complexes) and omit the second step (selection of complexes and main complexes from the candidates), because the computation time of the second step is negligible.

The total time complexity of searching for complexes depends on three factors, i.e., (1) computing the mutual information, (2) searching for MIPs, and (3) searching for complexes. The total computational cost is roughly bounded above by the products of these three factors (1)–(3). Each factor is given as follows.

1. The time complexity of computing the mutual information depends on the type of probability distribution. If the probability distribution is Gaussian, as we assume in experiments in Section 3, then the time complexity is *O*(*N* ^3^).
2. For searching for MIPs, we utilize Queyranne’s algorithm in this study. The time complexity of Queyranne’s algorithm is known to be *O*(*N* ^3^) [35].
3. HPC finds MIPs of *N* − 1 subsets as shown in Section 2.5. Thus, the time complexity of HPC is *O*(*N*).

Therefore, the total time complexity is about *O*(*N*^3^) × *O*(*N*^3^) × *O*(*N*) = *O*(*N*^7^). We evaluate actual computation time in Section 3.2 using a simulation.

### 2.6 A relation between monotonicity and submodularity

Before moving to the experiments, we show in this subsection a relation between monotonicity (Eq (13)) and submodularity (Def. 1). We show that monotonicity is satisfied by a broader class of submodular functions. Based on this relation, we show that we can extend our framework to the class of submodular functions.

As shown in Section 2.5.1, the mutual information has monotonicity, i.e., *I*(*T*_L_; *T*_R_) ≥ *I*(*S*_L_; *T*_R_) (Eq (13)). This monotonicity can be derived only from the submodularity of the entropy, as follows.

*Proof*.

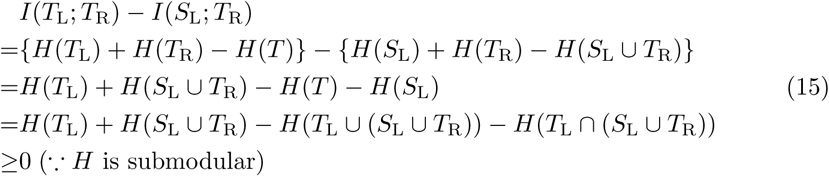

□

This relation between the submodularity of the entropy and the monotonicity of the mutual information is known as a relation between the submodularity of the entropy and the non-negativity of the conditional mutual information [21]. This proof indicates that the relation between the monotonicity and the submodularity can be stated in a general form [36]:

#### Proposition 9.

*Let f* : 2^*V*^ → ℝ *be a submodular function, and g* : 2^*V*^ × 2^*V*^ → ℝ *be a function defined as*

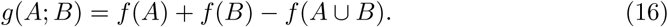

*Let T be a subset of V and* (*T*_L_, *T*_R_) *be a partition of T*. *Then, the function g satisfies the following inequality for any non-empty subset S*_L_ *of T*_L_ *(S*_L_ ⊂ *T*_L_).

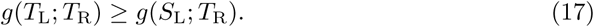

This proposition means that, given any submodular function *f*, we can define a new function *g* that satisfies the monotonicity. Consequently, the function *g* satisfies all the auxiliary theorems shown in Section 2.5.2, as the mutual information does. Therefore, if we regard *g* as an information loss function (although it does not have to be “informational”), and define complexes using *g*, we can use HPC to search for the complexes. Note that a function with the form Eq (16) satisfies the monotonicity, but not the other way around, i.e., a function that satisfies the monotonicity is not necessarily represented in the form of Eq (16).

#### Algorithm 1 Hierarchical Partitioning for Complex search (HPC)

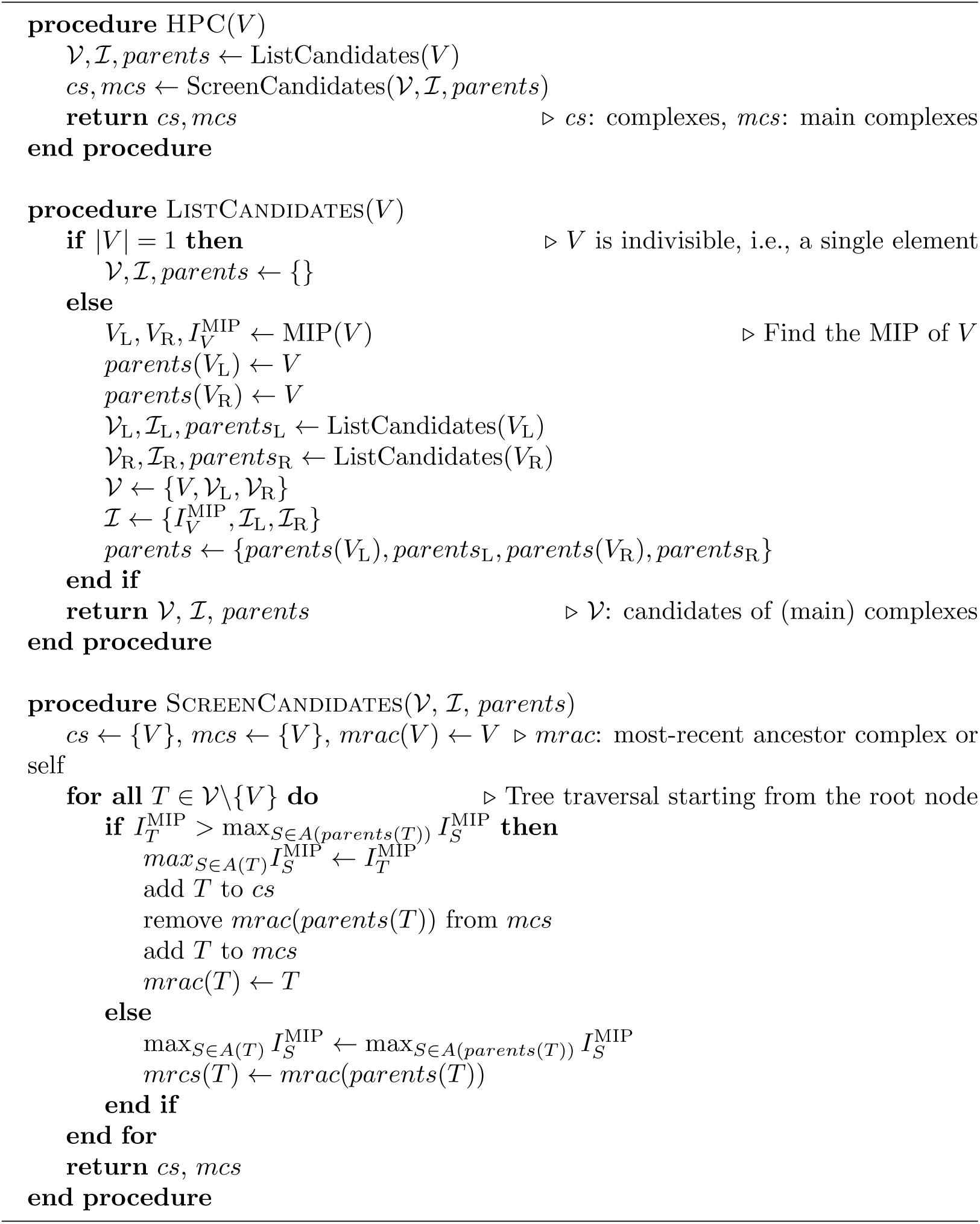

In addition, the function *g*(*S*_L_; *S*_R_) = *g*_*S*_ (*S*_L_) is a symmetrized submodular function (Section 2.3.2). This means that we can use Queyranne’s algorithm to search for MIPs when we use *g* as an information loss function.

Thus, if we regard a symmetrized submodular function as an information loss function, we can utilize Queyranne’s algorithm and HPC to search for MIPs and complexes, respectively. This is summarized as follows.

#### Submodular Complex (Complex for symmetrized submodular functions)

Given any submodular function *f* : 2^*V*^ → ℝ, we consider a function *g* defined as

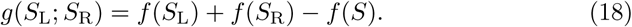

Then, the function *g* satisfies the same properties as the mutual information, i.e., symmetric-submodularity and monotonicity. Therefore, if we regard *g* as an information loss function, we can use Queyranne’s algorithm and HPC to search for MIPs and complexes, respectively.

## 3 Results

We first evaluated the performance of the proposed algorithm HPC using simulated data in Sections 3.1 and 3.2. We then applied HPC to a real neural dataset in Section 3.3. The MATLAB codes for reproducing the results in the experiments are available at https://github.com/oizumi-lab/PhiToolbox.

Throughout the simulations below, we consider a first-order autoregressive (AR) model with Gaussian noise,

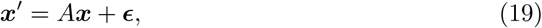

where ***x*** and ***x***′ are the present and the past states of a system consisting of *N* elements, *A* is an *N* × *N* matrix called the connectivity matrix, and ***ϵ*** is Gaussian noise with mean 0 and covariance Σ(***E***). We consider the stationary distribution of this AR model. The stationary distribution of *p*(***x***) is a Gaussian distribution with mean 0 and covariance Σ(***X***). The covariance Σ(***X***) can be computed from the following discrete Lyapunov equation

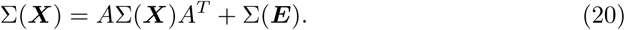

By using the covariance matrix Σ(***X***), the mutual information can be analytically calculated (see S2 Text). The details of the parameter settings are described in each subsection.

### 3.1 A simple example

We demonstrate how the proposed algorithm HPC works by considering a simple exemplary model. We consider an AR model with six elements (*N* = 6). The connectivity matrix *A* of the model is shown in Fig 5 as a network. This connectivity matrix *A* is symmetric and consists of two modules, one consisting of two elements, 1 and 2, and the second consisting of four elements, 3–6. The intra-connection strength in each group is 0.05*/N*, except the connection between the elements 5 and 6. The connection strength between 5 and 6 is 0.1*/N*, which is stronger than that between the other pairs. The inter-connections between the groups are 0.01*/N*, which are weaker than the intra-connections in each group. The strength of self connections is set to 0.9*/N*. The covariance Σ(***E***) of Gaussian noise is set to 0.01*I*, where *I* is an identity matrix.

**Fig 5.**
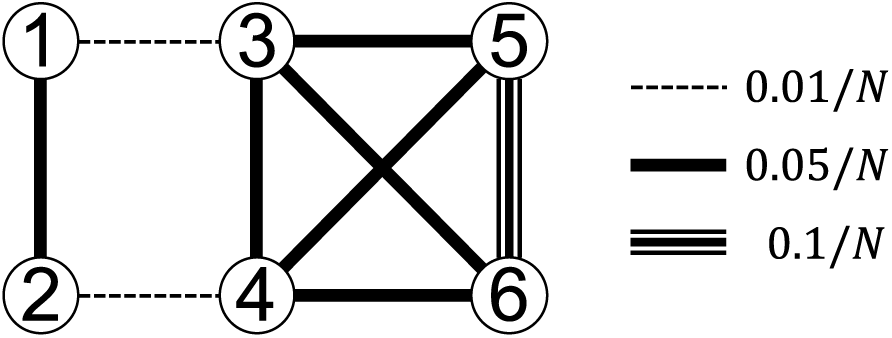
The connectivity matrix *A* of the AR model used in Section 3.1. The matrix *A* is symmetric, i.e. *a*_*ij*_ = *a*_*ji*_. The self connections (*a*_*ii*_) are omitted from this figure for simplicity.

Figure 6 shows how HPC hierarchically divides the system with MIPs. First, since the connections between two subsystems {1, 2} and {3, 4, 5, 6} are weak, the partition that splits the system into {1, 2} and {3, 4, 5, 6} is the MIP of the entire system. Then, these two subsystems are further divided with their MIPs. The first subsystem {1, 2} is divided into single elements {1} and {2}. The second subsystem {3, 4, 5, 6} is successively divided into {3} and {4, 5, 6}, {4} and {5, 6}, and {5} and {6}. Among these subsystems, {1, 2}, {3, 4, 5, 6} and {5, 6} are identified as complexes by comparing the amount of mutual information of the subsystems for their MIPs, as described in Section 2.5. Furthermore, the subsystems {1, 2} and {5, 6} are identified as main complexes.

**Fig 6.**
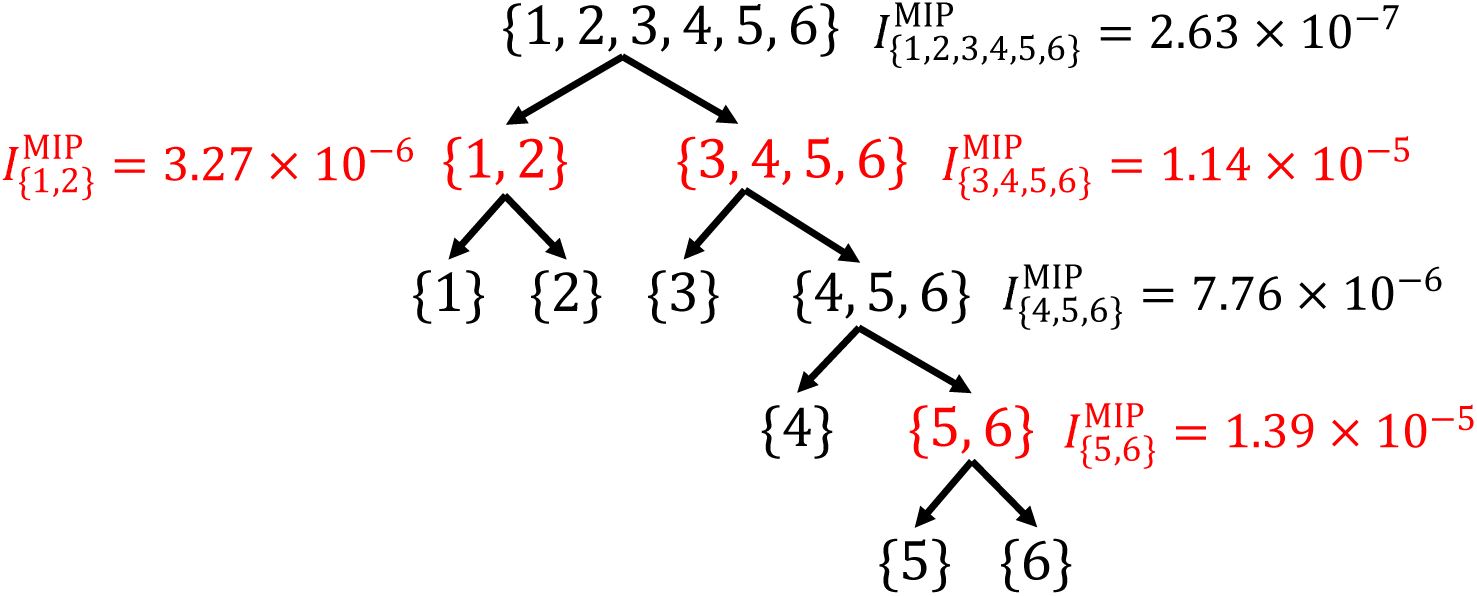
The way HPC divided the system at MIPs. First, the entire system was divided into two subsystems {1, 2} and {3, 4, 5, 6}. Then, the first subsystem {1, 2} was divided into single elements {1} and {2}. The second subsystem {3, 4, 5, 6} was successively divided into {3} and {4, 5, 6}, {4} and {5, 6}, and {5} and {6}. Among these subsystems, {1, 2}, {3, 4, 5, 6} and {5, 6} are complexes, which are indicated by red color. The subsystems {1, 2} and {5, 6} are main complexes.

Figure 7 visualizes the relation among complexes, main complexes, and subsets that are not complexes. For example, the subsystem {4, 5, 6} is not a complex, because it is at a lower position than one of its ancestors {3, 4, 5, 6} in the hierarchy. In contrast, the subsystem {3, 4, 5, 6} is a complex, because it is at higher position than its only ancestor {1, 2, 3, 4, 5, 6}. The subsystems {1, 2} and {5, 6} are main complexes because they are at the locally highest positions. HPC evaluates *I*^MIP^ of only five subsystems, while the exhaustive search evaluates that of 57(= 2^6^ − 7) subsystems. Thus, our HPC algorithm efficiently finds the complexes and the main complexes by hierarchically partitioning the system.

**Fig 7.**
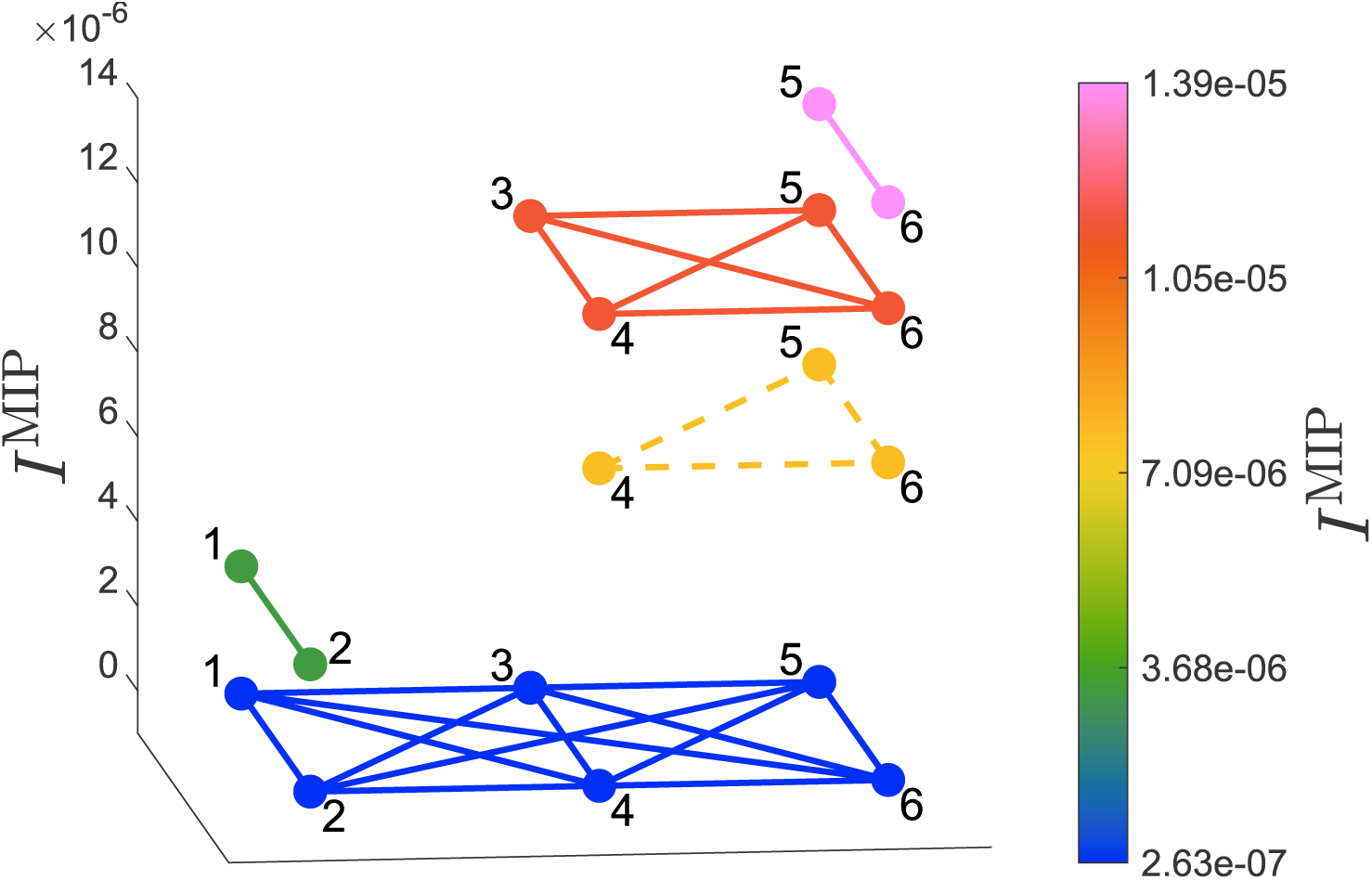
The amount of the mutual information *I*^MIP^ of subsystems appeared in the dividing process of HPC. The elements in a subsystem are connected by edges in the same color. The color indicates the amount of the mutual information *I*^MIP^. The subsystems with solid lines are (main) complexes, and that with dashed lines is not a complex.

### 3.2 Computation time of HPC

Next, we empirically measured the computation time of searching for complexes by simulation.

We randomly generated the connectivity matrices *A* in Eq (19). We determined each element of this connectivity matrix *A* by sampling from a Gaussian distribution with mean 0 and variance 0.01*/N*, where *N* is the number of elements. The covariance Σ(***E***) of the additive Gaussian noise in the AR model was set to 0.01*I*. All computation times were measured on a machine with an Intel Xeon Gold 6154 processor at 3.00GHz. All the calculations were implemented in MATLAB 2019a.

Figure 8 shows the log–log plot of the actual computation time. For comparison with the HPC algorithm, we also measured the computation time when complexes are exhaustively searched for by brute force. The red circles indicate the computation time of the proposed algorithm HPC. To estimate the order of computation time, we fitted a linear function to the red circles by the minimum mean squared error estimation. We discarded the first five circles (*N* ≤ 50) from the fitting analysis, because, when *N* is small, computation time is affected by lower-order terms. We obtained the red solid line, log_10_ *T* = 5.139 log_10_ *N* − 7.104, shown in Fig 8. As can be seen, the red solid line well approximates the red circles for larger *N*. This means that the computation time of HPC increases in polynomial order (*T* ∝ *N* ^5.139^). This is reasonably bounded above by the theoretical estimate of the computation time *T* ∝ *N* ^7^ (Section 2.5.4). The main reason why the empirical computation speed is faster than the theoretical estimate is thanks to the computation of the mutual information by MATLAB. As we mentioned in Section 2.5.4, the time complexity of the computation of mutual information for Gaussian distributions is *O*(*N* ^3^). However, the actual computation time grows more slowly (from *O*(*N*) to *O*(*N* ^2^)), when *N* is up to several hundred. In contrast, when we exhaustively searched for complexes, the computation time grows exponentially as indicated by the black triangles. We fitted an exponential function to the black triangles by the minimum mean square error estimation, and also discarded the first five points (*N* ≤ 7) from the fitting analysis. We obtained the black dashed curve, *T* ∝ 2.4689^*N*^ as shown in Fig 8.

**Fig 8.**
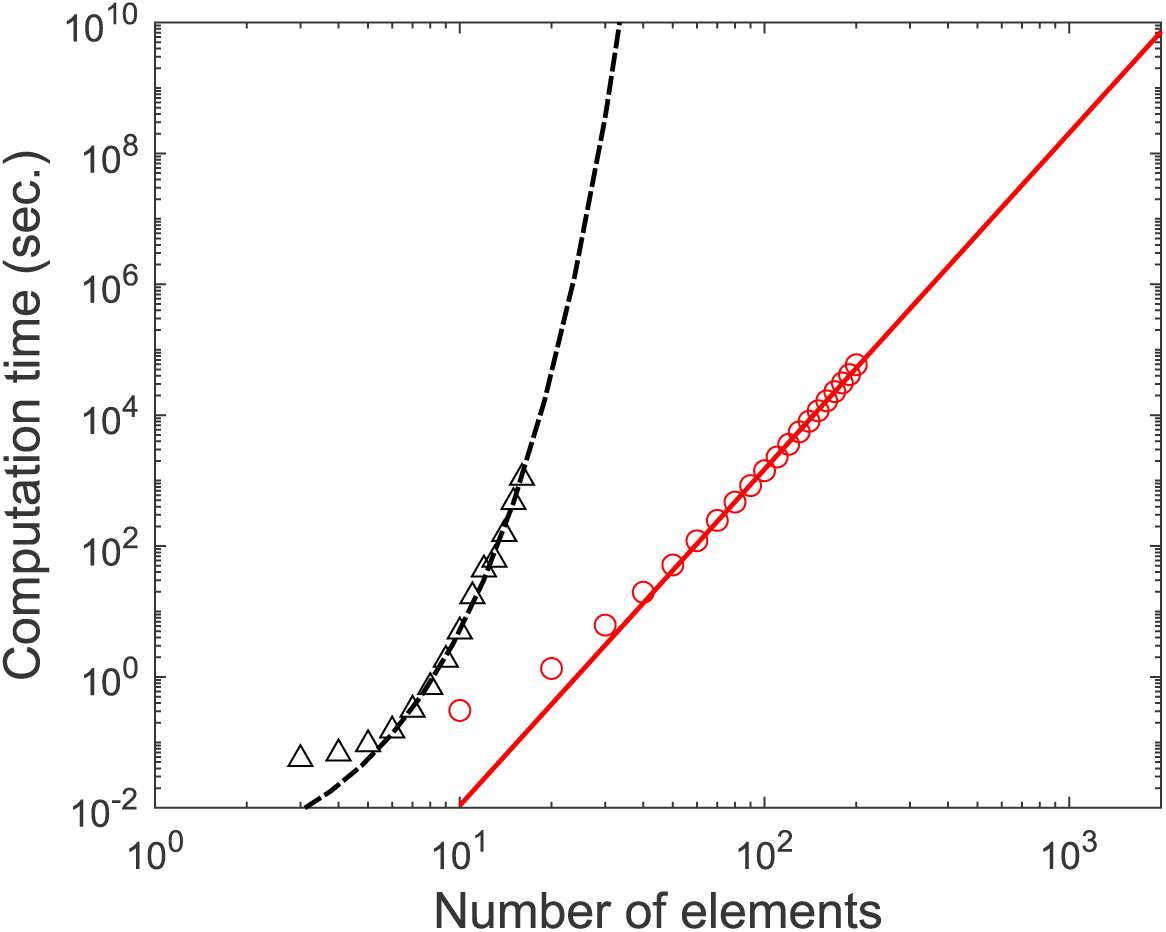
Computation time of the proposed algorithm HPC and the exhaustive search. The red circles and the red solid lines indicate the computation time of HPC and the fitted linear function (log_10_ *T* = 5.139 log_10_ *N* − 7.104). The black triangles and black dashed lines indicate the computation time of the exhaustive search and the fitted exponential function (log_10_ *T* = 0.3925*N* − 3.225).

As can be seen from Fig 8, HPC is much faster than the exhaustive search. For example, when *N* = 10^2^, the actual computation time of HPC was about 24 minutes, while that of the exhaustive search would be about 3.36 × 10^28^ years.

### 3.3 An application to real neural data

Finally, we applied HPC to a neural dataset to demonstrate how HPC can be used in real settings. We used electrocorticography (ECoG) data recorded from a macaque monkey. The dataset is available at an open database, Neurotycho.org (http://neurotycho.org/) [23]. One hundred twenty-eight electrodes were implanted in the left hemisphere. The electrodes were placed at 5-mm intervals covering the frontal, parietal, temporal and occipital lobes, and the medial frontal and parietal walls. Signals were sampled at a rate of 1 kHz. The monkey was awake with the eyes covered by an eye-mask to restrain visual responses. To remove line noise and artifacts, we performed bipolar re-referencing between adjacent electrode pairs, i.e. subtracting the signal of one electrode from that of the other. The number of bipolar re-referenced electrodes was 64 in total. Among the 64 channels, two channels were removed from further analysis because of measurement noise.

We extracted 15-minute signals and divided them into 1-minute time windows. Each 1-minute time window consists of 1 kHz × 60 s = 60,000 samples. We searched for complexes in each time window. We approximated the probability distribution of the signals with multivariate Gaussian distributions. Under the Gaussian approximation, we could compute the mutual information using the equation shown in S2 Text.

Figure 9 shows the main complexes and complexes in the first time window. In each panel, the main complexes and complexes are superimposed in ascending order of the amount of mutual information *I*^MIP^. We can see that the main complexes consist of pairs of nearby channels. In contrast, the complexes other than the main complexes, by definition, consist of more channels than the main complexes. In particular, the complexes with large *I*^MIP^ tend to consist of channels clustered in posterior areas: the complex with the smallest *I*^MIP^ among all complexes is the whole system (all 62 channels, shown in blue color). Then, the complexes with moderate *I*^MIP^ consist of channels clustered in the middle and posterior areas (shown in bluish-green color), and complexes with large *I*^MIP^ consist of channels clustered in posterior areas (shown in yellowish-green color). Thus, we can see that the complexes tend to consist of channels in the more posterior area as *I*^MIP^ increases. Every (main) complex at the first time window is separately shown in S1 Fig. We obtained a similar tendency at the different time windows: main complexes consist of pairs or triples of channels, and complexes cover a more posterior area as *I*^MIP^ increases (S2 Fig and S3 Fig).

**Fig 9.**
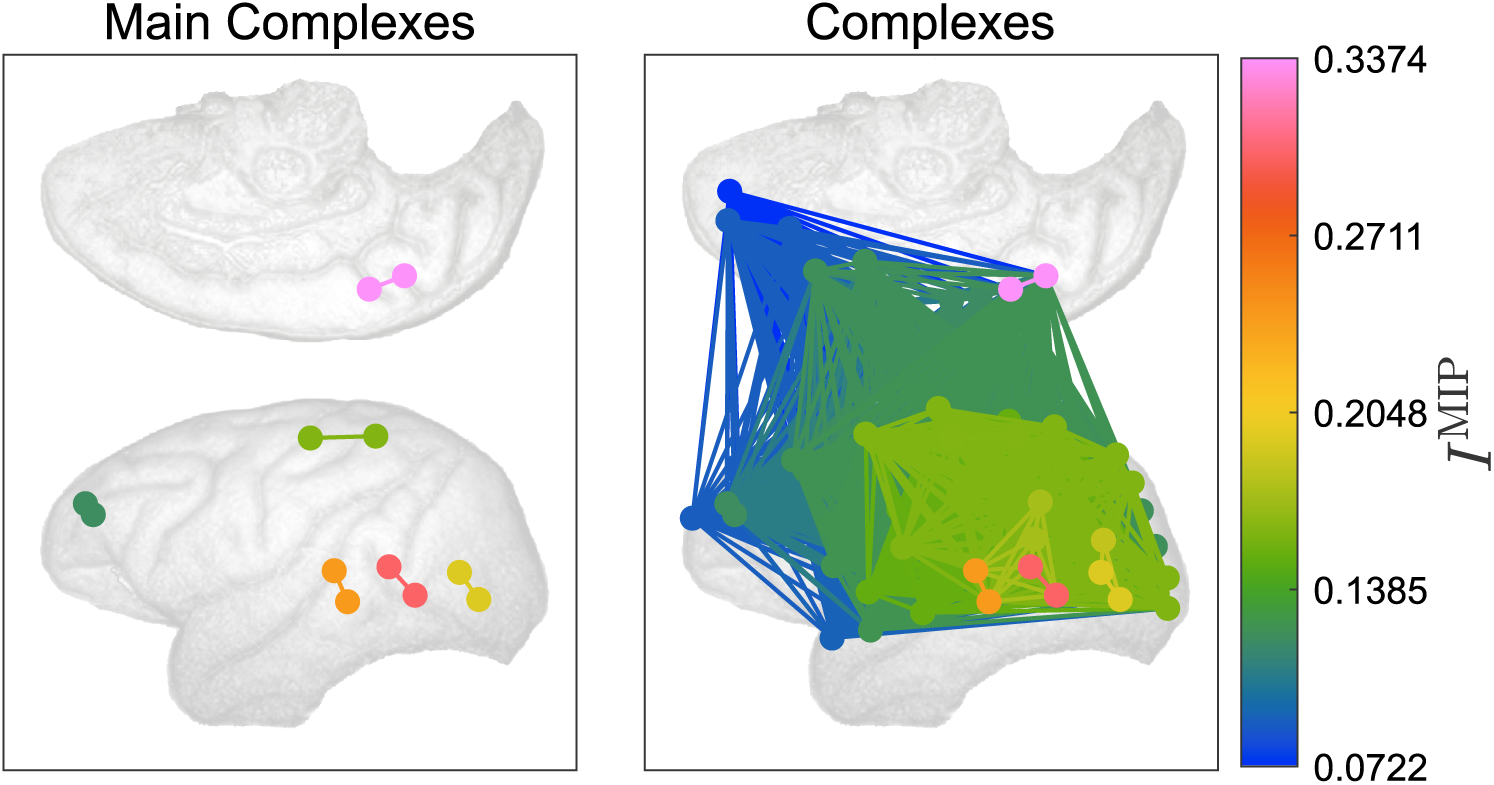
Main complexes and complexes at the first time window. The channels in each (main) complex are connected by edges with the same color. The color indicates the amount of the mutual information *I*^MIP^ of the (main) complex. The main complexes and complexes are superimposed in ascending order of the amount of mutual information *I*^MIP^.

Thus, in this experiment, since main complexes consist of only pairs or triples of channels, it seems difficult to gain insights from the main complexes only. It would appear better to analyze not only the main complexes but also these other complexes to extract meaningful information.

## 4 Discussion

In this study, we proposed a fast algorithm for searching for informational cores, or complexes. We call the proposed algorithm Hierarchical Partitioning for Complex search (HPC) because it narrows down candidates for complexes by hierarchically partitioning a system. We proved that the HPC algorithm can efficiently enumerate all the complexes and the main complexes when mutual information is used as an information loss function. The number of subsystems whose MIPs are evaluated in HPC is a linear order of system size. This is dramatically smaller than that of an exhaustive search, which is exponential of system size. We also proved that if we regard a specific type of symmetric submodular function, which we call a symmetrized submodular function, as an information loss function, we can apply HPC to search for complexes. In the experiments, we numerically evaluated the computation time of the overall complex search process by HPC and showed that the computation time was approximately *O*(*N* ^5^). We applied HPC to a simple model and monkey ECoG data to demonstrate how HPC can be applied.

In the monkey EcoG data analyses in Section 3.3, the main complexes were small and localized, i.e., pairs or triples of channels. This may be because the channels in a main complex were spatially so close to each other that the mutual information between them was high. In contrast, the complexes were larger than the main complexes, and there was a stable tendency for the complexes to cover the posterior area. Therefore, it appears important to consider not only main complexes but also complexes to uncover global characteristics of the brain network.

HPC enables us to search for (main) complexes in systems consisting of several hundred elements (*N* ∼ 100–200) in a practical amount of time. The number of channels in EEG or ECoG is typically within this range. The concept of (main) complex was originally proposed as a locus of consciousness in the integrated information theory of consciousness. However, it can be utilized to analyze brain networks in contexts unrelated to consciousness, and also to other probabilistic systems apart from the brain. HPC will therefore be beneficial not only for consciousness studies but also in other general research fields.

As shown in Section 2.5, the inequality in Proposition 5 is a prerequisite for HPC to be exact. While the mutual information satisfies the inequality, other information loss functions, e.g., stochastic interaction [37, 38], integrated information based on mismatched decoding [31], and geometric integrated information [25], do not necessarily satisfy it. Therefore, when these functions are utilized, there is no guarantee that HPC can find all the (main) complexes. Nevertheless, HPC might practically work well and might be used as an approximate algorithm. It would be interesting to test the extent to which HPC works well when these functions are utilized.

As shown in Section 2.6, our framework can be naturally extended by regarding a symmetrized submodular function as an information loss function. An example of such symmetrized submodular functions other than the mutual information is the weight of a graph cut. Since graph representations of systems are useful and important in many scientific fields, we will extend our framework to graphs in the next study.

## Supporting information

**S1 Text. Select complexes and main complexes from the candidate set** *V*

**S2 Text. Analytical formula of mutual information for Gaussian distribution**

**S1 Fig. The (main) complexes at the first time window in ECoG data analyses**. The (main) complexes at the first time window are sorted by the amount of mutual information *I*^MIP^ in descending order. The channels in each complex are connected by edges. The color of the markers and the edges indicates the amount of mutual information *I*^MIP^. The panels with a red box outline show main complexes and those with a black box outline show complexes that are not main complexes. We can see that the main complexes are small; in contrast, the complexes are larger than the main complexes; and the complexes tend to consist of channels in the more posterior area as *I*^MIP^ increases.

**S2 Fig. Main complexes at all 15 time windows**. The channels in each main complex are connected by edges with the same color. The color of the markers and the edges indicates the amount of mutual information *I*^MIP^. We can see that the main complexes are pairs or triples of channels.

**S3 Fig. Complexes at all 15 time windows**. The channels in each (main) complex are connected by edges with the same color. The color of the markers and the edges indicates the amount of mutual information *I*^MIP^. The main complexes and complexes are superimposed in ascending order of the amount of mutual information *I*^MIP^. The complexes at different time windows are not exactly the same, but we can see that the complexes tend to consist of channels in the more posterior area as *I*^MIP^ increases.

## Acknowledgments

This work was partially supported by JST CREST Grant Numbers JPMJCR15E2 including AIP challenge program and JPMJCR1864, and JSPS KAKENHI Grant Number 18H02713, Japan.

## Notes

https://github.com/oizumi-lab/PhiToolbox

### Summary of Updates

Supplementary material has been added and the image resolution has been raised because for some reason supplementary material was missing and the resolution became low.

