## Supplementary material for "Efficient search for informational cores in complex systems: Application to brain networks": S1 Text

### Supporting information

#### S1 Text. Select complexes and main complexes from the candidate set $\mathcal{V}$

In this subsection, we explain how we select complexes and main complexes from the candidate set $\mathcal{V}$, which is defined in Section 2.5.3.

##### Part A: Complex

We need to check whether each complex candidate belonging to $\mathcal{V}$ actually satisfies the condition in Def. 2. The condition means that a candidate must have larger $I^{\mathrm{MIP}}$ than its supersets to be a complex. To check this, the tree structure obtained by the hierarchical partitioning procedure can be utilized: the candidate needs to be compared only with its ancestor candidates in the tree because its supersets other than the ancestors straddle at least one of the MIP boundaries of the ancestors, and therefore have smaller $I^{\mathrm{MIP}}$ than the ancestors (Proposition 5). Thus, a candidate is a complex if it has larger $I^{\mathrm{MIP}}$ than its ancestor candidates. This comparison can be efficiently done by propagating the maximum value of $I^{\mathrm{MIP}}$ among ancestors and self from the root node (the whole system) to the leaf nodes (the single elements) as described below.

Here, let $A(\cdot)$ denote ancestors and self. For example, $A(V_{\mathrm{LLR}})$ indicates $\{V, V_{\mathrm{L}}, V_{\mathrm{LL}}, V_{\mathrm{LLR}}\}$. Starting from the root node $V$, the amount of mutual information $I_T^{\mathrm{MIP}}$ of each candidate $T \in \mathcal{V}$ is compared with the maximum of the mutual information of $T$'s ancestors $\max_{S \in A(parents(T))} I^{\mathrm{MIP}}(S)$:

$$\text{if } I_T^{\mathrm{MIP}} > \max_{S \in A(parents(T))} I_S^{\mathrm{MIP}}, \text{ then } \begin{cases} T \text{ is a complex, and} \\ \max_{S \in A(T)} I_S^{\mathrm{MIP}} = I_T^{\mathrm{MIP}}, \end{cases}$$

$$\text{otherwise } \begin{cases} T \text{ is not a complex, and} \\ \max_{S \in A(T)} I_S^{\mathrm{MIP}} = \max_{S \in A(parents(T))} I_S^{\mathrm{MIP}}. \end{cases}$$

By repeating this comparison while traversing the entire tree from the root to leaf nodes (i.e., in pre-order), we can list complexes without omission. In total, the comparison is done $|\mathcal{V}|$ times, where $|\mathcal{V}|$ indicates the number of subsystems in $\mathcal{V}$. We can easily show that $|\mathcal{V}| = N - 1$ as follows. Before the dividing steps, there is only one system (the whole system $V$) in $\mathcal{V}$. Then, two subsystems are added to $\mathcal{V}$ every dividing step. Since HPC divides the system $N - 1$ times as described above, there are $1 + 2(N - 1)$ subsystems in $\mathcal{V}$ at the end. Note, however, this number counts $N$ single elements. Therefore, we finally get $|\mathcal{V}| = 1 + 2(N - 1) - N = N - 1$.

##### Part B: Main complex

As stated in Section 2.5.3, a main complex is a locally farthest complex from the root in the tree. Based on this, we can easily find main complexes during the tree traversal as follows. If we find a complex, we temporarily add it to the list of main complexes. Then, if we find another complex that is a descendant of a temporal main complex, we remove the temporal main complex from the list and add the new one to the list. On finishing the tree traversal, we have obtained a complete list of main complexes.
