## Supplementary material for "Efficient search for informational cores in complex systems: Application to brain networks": S2 Text

### Supporting information

#### S2 Text. Analytical formula of mutual information for Gaussian distribution

We describe the analytical formula of the mutual information when the probability distribution is Gaussian. Let us begin by introducing the notation. We consider a probabilistic system consisting of $N$ elements. We represent the states of the system as $\boldsymbol{x}_S = (x_1, \ldots, x_N)$. We assume that the probability distribution $p(\boldsymbol{x}_S)$ is Gaussian:

$$\begin{aligned} p(\boldsymbol{x}_S) &= \frac{1}{\sqrt{|2\pi\Sigma|}} \exp\left(-\frac{1}{2}\boldsymbol{x}_S^T \Sigma^{-1} \boldsymbol{x}_S\right), \\ &= \frac{1}{\sqrt{|2\pi\Sigma|}} \exp\left(-\frac{1}{2}(\boldsymbol{x}_{S_\mathrm{L}}^T, \boldsymbol{x}_{S_\mathrm{R}}^T) \begin{pmatrix} \Sigma_\mathrm{L} & \Sigma_\mathrm{LR} \\ \Sigma_\mathrm{LR}^T & \Sigma_\mathrm{R} \end{pmatrix}^{-1} \begin{pmatrix} \boldsymbol{x}_{S_\mathrm{L}} \\ \boldsymbol{x}_{S_\mathrm{R}} \end{pmatrix}\right), \end{aligned}$$

where $\Sigma$, $\Sigma_\mathrm{L}$ $\Sigma_\mathrm{R}$ are the covariance matrices of $\boldsymbol{x}_S$, $\boldsymbol{x}_{S_\mathrm{L}}$, and $\boldsymbol{x}_{S_\mathrm{R}}$, $\Sigma_\mathrm{LR}$ is the covariance matrix between $\boldsymbol{x}_{S_\mathrm{L}}$, and $\boldsymbol{x}_{S_\mathrm{R}}$, and $|\cdot|$ indicates the determinant. Then, we get the marginalized distributions:

$$\begin{aligned} p(\boldsymbol{x}_{S_\mathrm{L}}) &= \frac{1}{\sqrt{|2\pi\Sigma_\mathrm{L}|}} \exp\left(-\frac{1}{2}\boldsymbol{x}_{S_\mathrm{L}}^T \Sigma_\mathrm{L}^{-1} \boldsymbol{x}_{S_\mathrm{L}}\right), \\ p(\boldsymbol{x}_{S_\mathrm{R}}) &= \frac{1}{\sqrt{|2\pi\Sigma_\mathrm{R}|}} \exp\left(-\frac{1}{2}\boldsymbol{x}_{S_\mathrm{R}}^T \Sigma_\mathrm{R}^{-1} \boldsymbol{x}_{S_\mathrm{R}}\right). \end{aligned}$$

Note that we can assume the mean of the Gaussian distribution is zero without loss of generality because the mean value does not affect the values of mutual information.

The entropy of an $N$-dimensional Gaussian distribution with a covariance matrix $\Sigma$ is given by $\frac{1}{2}\log|2\pi\Sigma| + \frac{N}{2}$. By substituting this expression into Eq. (5), we get

$$I(S_\mathrm{L}; S_\mathrm{R}) = \frac{1}{2}\log\frac{|\Sigma_\mathrm{L}||\Sigma_\mathrm{R}|}{|\Sigma|}.$$
