## Supplementary figures and images for "Efficient search for informational cores in complex systems: Application to brain networks"

### S1 Fig

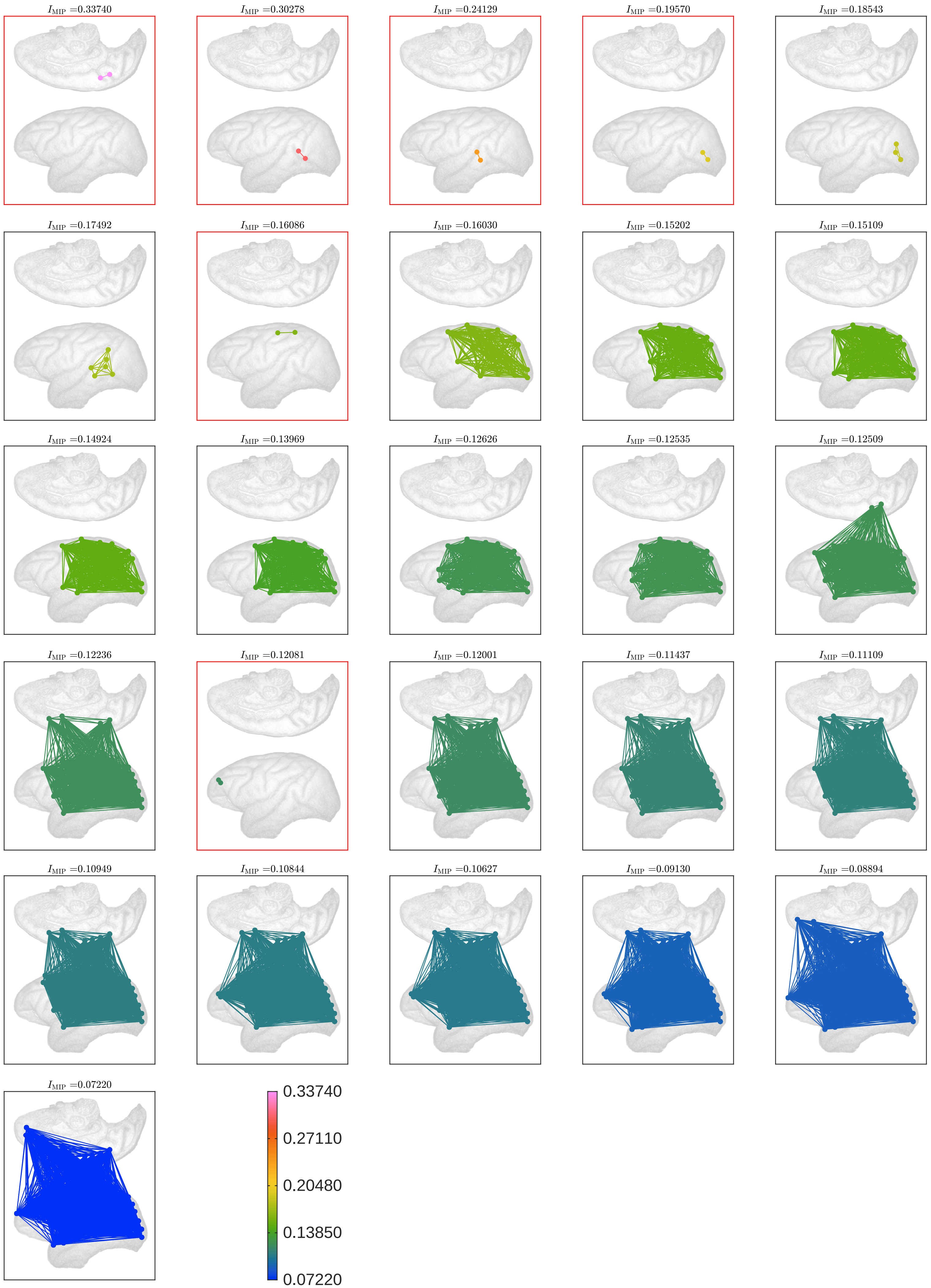

### S2 Fig

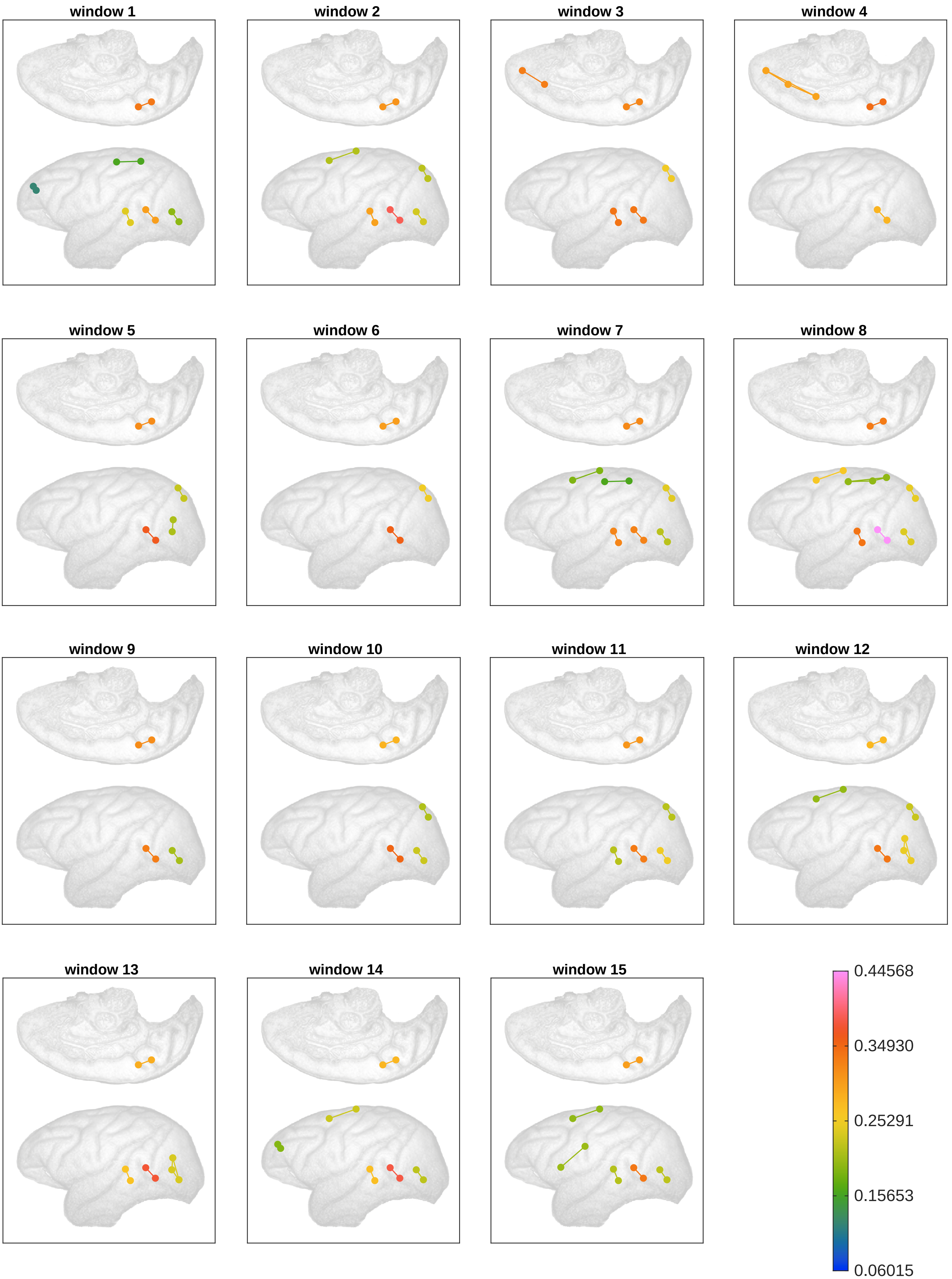

### S3 Fig

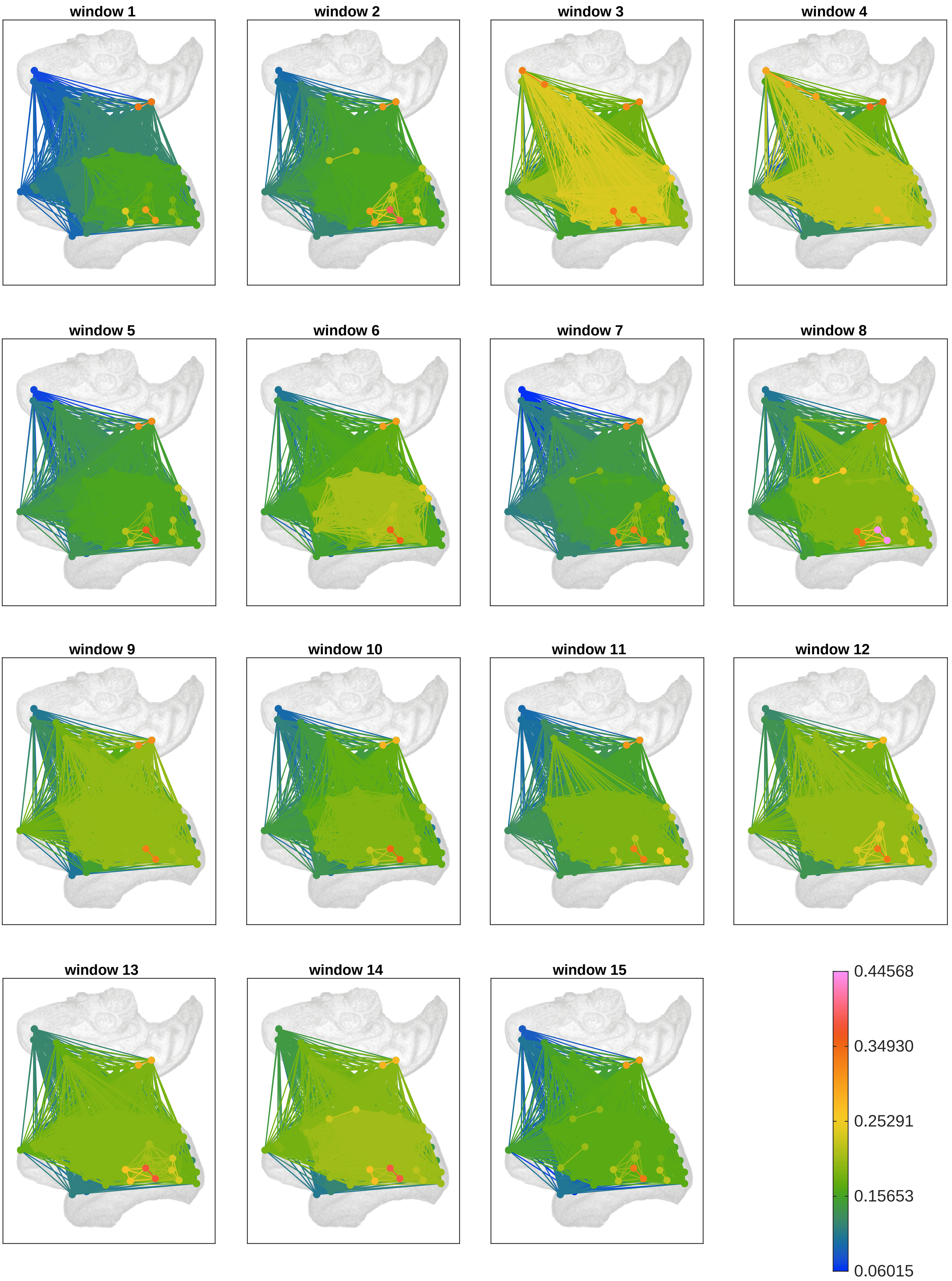
